# Sustained neurotrophic factor co-treatment enhances donor and host retinal ganglion cell survival in mice

**DOI:** 10.1101/2024.03.07.583961

**Authors:** Jonathan R. Soucy, Julia Oswald, Emil Kriukov, Monichan Phay, John Masland, Christian Pernstich, Petr Baranov

**Author notes:** Co-first authors (Jonathan R. Soucy, Julia Oswald).

## Abstract

Retinal ganglion cells (RGCs) lack regenerative capacity in mammals, and their degeneration in glaucoma leads to irreversible blindness. Autologous and allogeneic RGC replacement with stem cell-derived neurons has been established as a promising strategy for vision restoration, but it has been limited by relatively low (<1%) survival rates of donor cells in the hostile microenvironment of a diseased retina and optic nerve. Brain-derived neurotrophic factor (BDNF) and glial-derived neurotrophic factor (GDNF) are known to support the survival of neurons, including RGCs in. vitro and in vivo. Here, we aim to improve donor RGC survival by supplementing our in vitro cultures and in transplants with a slow-release formulation of these neuroprotective agents. We show that slow-release BDNF/GDNF significantly enhances RGC differentiation, survival, and function in vitro. Furthermore, we demonstrated that BDNF/GDNF co-treatment improved mouse and human stem cell-derived donor RGC transplantation outcomes by 2.7- and 15-fold in mice, respectively. Lastly, we show that slow-release BDNF/GDNF provides neuroprotective effects on host RGCs, preserving retinal function in a model of optic neuropathy. Altogether, this approach of engineering the retinal microenvironment with slow-release neurotrophic factors significantly enhances both donor and host neuron survival, representing a promising approach for treating glaucoma and other optic neuropathies.

**Table of Contents Graphic:** 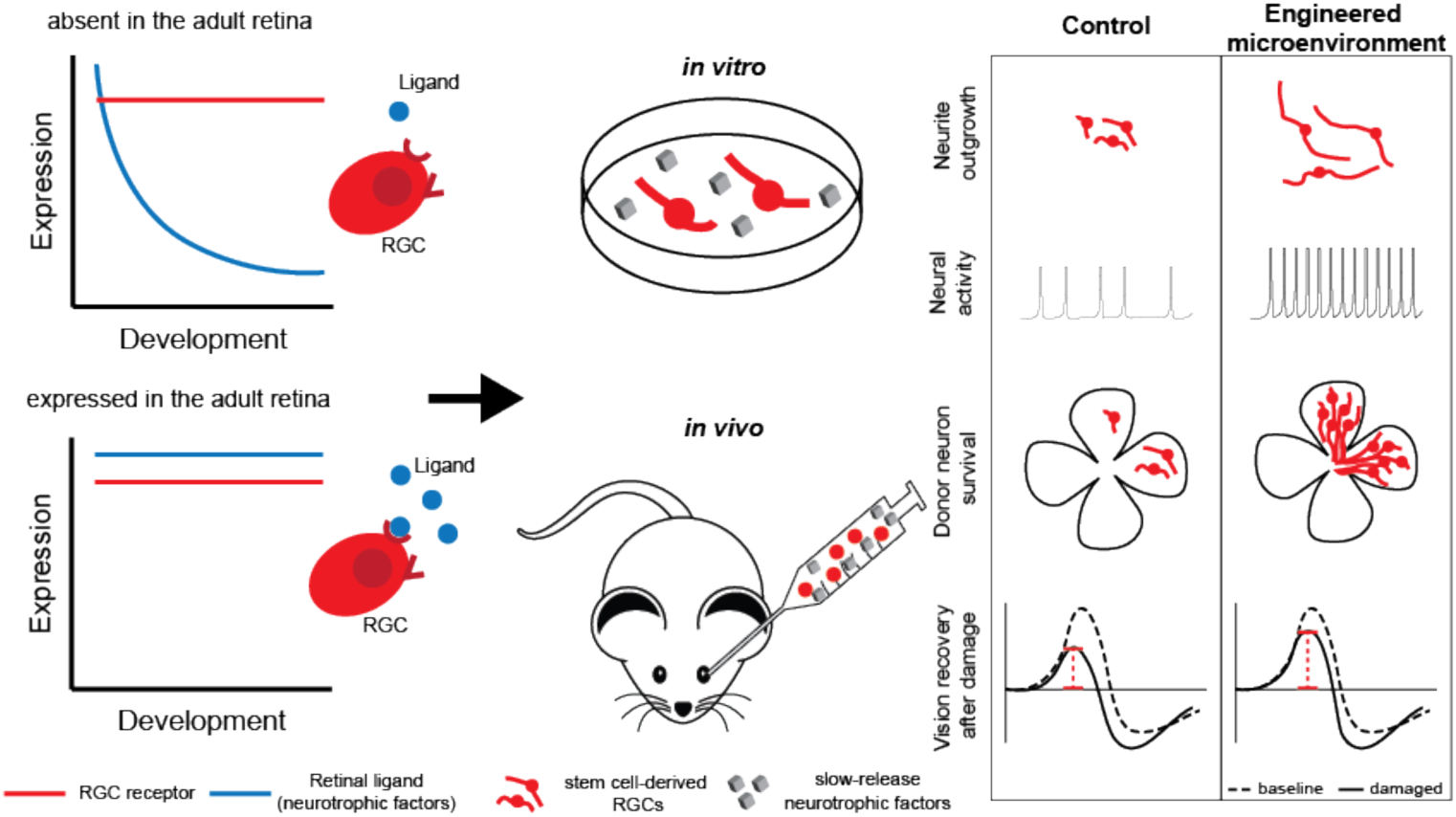

Engineering the retinal environment with neurotrophic factors to mimic development improves donor and host neuron survival, outgrowth, and function in an optic neuropathy mouse model.

## Introduction

The loss or degeneration of retinal ganglion cells (RGCs) in glaucoma is a leading cause of irreversible blindness worldwide, with no current therapies available to restore vision once these cells are lost.^1^ While RGCs, the neurons in the retina that relay all visual information from the eye to the brain, can regenerate in some lower vertebrates, they have no capacity for regeneration in the mammalian retina.^2^ Cell transplantation has been proposed as a potential solution to replace RGCs lost in advanced retinal disease, but low cell survival rates post-transplantation have limited the effectiveness and further development of this strategy.^3–6^

While RGC transplantation has been studied using both primary and stem cell-derived neurons in different pre-clinical models, each approach has distinct benefits and limitations.^3,6^ Xenogeneic transplantation uniquely enables the study of human RGCs within rodent models, significantly enhancing clinical relevance and informing translational strategies, yet involves heightened immune rejection challenges. Primary RGC transplants from donor retinas offer proven integration but are constrained by limited cell availability and scalability.^4^ These limitations have led to the development of stem cell-derived RGCs, including those from human embryonic stem cells (hESCs) and induced pluripotent stem cells (iPSCs), to facilitate large-scale and defined cell production. However, achieving optimal donor cell survival, maturation, and integration within the host retinas remains an ongoing research challenge.

The glaucomatous retina is typically characterized by increased intraocular pressure, which can cause inflammation, immune cell infiltration, and host cell reactivity.^7,8^ Even after the intraocular pressure normalizes, the altered microenvironment with hyperactive glia can persist,^8^ making it challenging for transplanted donor cells to survive in the host retina. Further, after intravitreal transplantation, donor cells are assumed to encounter an environment in the vitreous cavity that lacks adequate oxygenation and metabolic support until they structurally integrate into their respective positions within the host retina.^6,9^ This deficiency can intensify the stress experienced by the donor cells, leading to further decreased survival rates.

Brain-derived neurotrophic factor (BDNF) and glial-derived neurotrophic factor (GDNF) are known to support host retinal neuroprotection in vivo and improve neuron survival in vitro.^6,10–16^ Moreover, since RGCs are one of the primary sources of BDNF and GDNF in the retina,^11,17,18^ their loss in diseases like glaucoma may create an environment that lacks these crucial survival factors. Therefore, supplementing transplanted cells with BDNF and GDNF could not only enhance the survival of these cells but also potentially rejuvenate the surrounding retinal tissue, creating a more favorable environment for transplantation, regeneration, and repair.

In this report, we differentiate both mouse and human stem cell-derived RGCs using a 3D retinal organoid approach. To improve the RGC differentiation yield and survival, we first investigated how supplementing the culture media with slow-release BDNF and GDNF can affect our retinal organoid cultures. We then studied RGC structural and functional characteristics in vitro and gene expression in silico to characterize donor cell response to slow-release BDNF and GDNF, showing increased neurite outgrowth and neural activity and reduced apoptotic gene expression. To make the retinal microenvironment more hospitable for transplantation and improve donor RGC survival in vivo, we transplanted both mouse and human stem cell-derived RGCs with slow-release formulations of BDNF and GDNF into the mouse eye. By quantifying these transplants using a semi-automated approach,^19^ we confirm that co-delivery of mouse and human stem cell-derived RGCs with slow-release formulations of BDNF and GDNF improves donor RGC coverage within the retina and the total number of cells that survive post-transplantation. After transplantation, we also investigated whether there was a recovery in visual function using an electroretinogram (ERG). While there was no recovery after transplantation with donor RGCs alone, we demonstrated that a slow-release formulation of BDNF and GDNF provides a neuroprotective effect on host RGC, preventing a decrease in ERG-related response caused by glutamate-induced neurotoxicity. Altogether, here we demonstrated the potential for microenvironment engineering using slow-release neurotrophic factors for both in vitro and in vivo applications.

## Materials and Methods

### Mouse Stem Cell-derived RGC Differentiation

Mouse stem cell-derived RGCs were differentiated using a 3D retinal organoid approach as previously described.^5,19^ Induced pluripotent stem cells derived from Tg[Thy1.2-Green Fluorescent Protein (GFP)]M mice were cultured on gelatin-coated (0.2% v/v) flasks in standard culture conditions (37°C, >95% humidity, 5% CO2). Mouse stem cell maintenance medium [10% fetal bovine serum, 1x L-glutamine, 1x non-essential amino acids, 1x sodium pyruvate, 1x antibiotic-antimycotic, 1x EmbryoMax Nucleosides, 100 μM β-Mercaptoethanol, and 100 U/mL mouse leukemia inhibitory factor in DMEM/F12 (3500 mg/L dextrose)] was replaced every two days until the stem cell colonies approached approximately 70% confluency. Upon reaching 70% confluency, stem cells were harvested using 1x trypsin-EDTA and either transferred to new gelatin-coated flasks at a density of 2000 cells/cm² or used for RGC differentiation as previously detailed. For RGC differentiation, 1500 cells were initially seeded in 50 μL of Optic Vesicle medium [1x L-glutamine, 1x non-essential amino acids, 1x sodium pyruvate, 1x antibiotic-antimycotic, 1x lipid concentrate, 0.2x Insulin-Transferrin-Selenium-Ethanolamine, 100 μM β-Mercaptoethanol, 1.5% fetal bovine serum, 1 μM N-Acetyl-L-cysteine, and 10 μM forskolin in DMEM/F12 (3500 mg/L dextrose)] per well in uncoated 96-well U-bottom polystyrene plates. After 24 hours in standard culture conditions, each well received an additional 50 μL of Optic Vesicle medium with 2% Matrigel to start forebrain/retinal differentiation. The next day, 100 μL of Optic Vesicle medium with 1% Matrigel was added to each well. The organoids were maintained in this medium until the ninth day of differentiation, with half of the medium being replaced every two days. On day 8, the organoids were extracted from the plates, randomly sectioned to isolate optic vesicles for expanded growth, and transferred to low-attachment flasks. From this point, they were kept in Optic Cup medium [1x L-glutamine, 1x non-essential amino acids, 1x sodium pyruvate, 1x antibiotic-antimycotic, 1x lipid concentrate, 100 μM β-Mercaptoethanol, 1x B27 without vitamin A, 1 μM N-Acetyl-L-cysteine, and 10 μM forskolin in DMEM/F12 (3500 mg/L dextrose)]. Half the medium was replaced every other day until days 21 to 24 of retinal maturation. Starting from day 16, the Optic Cup medium was supplemented with 250nM all-trans-retinoic acid.

### Human Stem Cell-derived RGC Differentiation

Colonies of H9-BRN3B:tdTomatoThy1.2-hESCs were cultured on six-well plates coated with 2% Matrigel with mTeSR1 media in a low oxygen incubator (5% CO2, 3.5% O2, above 95% humidity, 37°C). Media was replaced daily. When the cell density reached around 70% after 4-5 days, the cells were detached using accutase for 8 minutes at room temperature and transferred to a new Matrigel-coated plate at a ratio between 1:10 and 1:20.

Human RGCs were differentiated in 3D retinal organoid cultures as previously described.^19–21^ During cell passaging, hESC colonies were enzymatically dissociated for 12 minutes to form a single-cell suspension. To create embryoid bodies, 7,500 hESCs per well were seeded in an ultra-low attachment 96-well U-bottom plate in a low oxygen incubator for 24 hours. After 24 hours, the plate was moved to standard culture conditions. The differentiation media was exchanged daily and transitioned from a 3:1 mix of mTeSR1 and BE6.2-NIM [<u>B</u>27 + <u>E6</u> at <u>2</u>X concentration – neural induction medium: 1x L-glutamine, 1x sodium pyruvate, 1x antibiotic-antimycotic, 1x B27 without Vitamin A, 0.88 mg/mL NaCl, 38.8 mg/L insulin, 128 mg/L L-ascorbic acid, 28 μg/L selenium, 21.4 mg/L transferrin, and 38.8 mg/L NaHCO_3_ in DMEM/F12 (3500 mg/L dextrose)] to primarily BE6.2-NIM by day 8. To support neural and early retinal development, the media was supplemented with 20 μM Y-27632 on day 0, 1% Matrigel from days 2 to 6, 3μM IWR-1-endo from days 1 to 4, and 55 ng/mL BMP4 on day 6. On day 8, organoids were randomly sectioned to isolate optic vesicles, facilitating growth and keeping size under the diffusion limit (1.4 mm diameter). These chopped organoids were then placed on tissue culture-treated plastic in Long-Term Retinal Differentiation Media (LT-RDM) [1x L-glutamine, 1x sodium pyruvate, 1x antibiotic-antimycotic, 1x B27, 1x non-essential amino acids, and 10% fetal bovine serum in 50:50 1x DMEM/F12 (3500 mg/L Dextrose):DMEM (4500 mg/mL dextrose)] to encourage attachment and retinal epithelium growth. The media was replaced every two days, transitioning to a 1:1 mix of LT-RDM and Retinal Differentiation Media (RDM) [LT-RDM without fetal bovine serum] by day 17, with added 100nM smoothened agonist (SAG) from days 12 to 16. On day 17, the retinal tissue clusters were detached and moved to a poly(2-hydroxyethyl methacrylate)-coated flask for suspension culture in RDM supplemented with 100nM SAG. From day 18 onwards, the media, now a 1:3 mix of LT-RDM and RDM with 2.5% fetal bovine serum and 100nM SAG, was replaced every other day, gradually transitioning to pure LT-RDM by day 24. From day 26, the media was supplemented with 1mM taurine, and from day 32, 250nM all-trans-retinoic acid. The LT-RDM was replaced every other day until retinal organoids were collected for dissociation.

### RGC Isolation

To dissociate mature mouse (d21 – 24) and human retinal organoids (d44 – 48), the embryoid body dissociation kit from Miltenyi Biotec (catalog number 130-096-348) was used with the gentleMACS™ Dissociator, following the instructions provided by the manufacturer, with a single adjustment. The preset gentleMACS programs EB_01 and EB_02 were substituted with Brain_01 and Brain_02, respectively. After dissociation, mouse and human RGCs (Thy1.2-positive cells) were isolated by magnetic microbead-based sorting using the Dynabeads™ FlowComp™ Mouse Pan T (CD90.2) Kit from Invitrogen (catalog number 11465D), following the manufacturer’s instructions.

### Flow Cytometry

Dissociated cells were resuspended in 4% paraformaldehyde and fixed for 30 minutes on ice. Post-fixation, cells were centrifuged at 300 x g for 5 minutes and resuspended in isolation buffer [2 mM EDTA, 2% fetal bovine serum, and 0.1% bovine serum albumin in Hank’s Balanced Salt Solution without Ca2+ and Mg2+] for flow cytometric analysis. The samples were analyzed using a MACSQuant Analyzer 10 flow cytometer. The data were gated based on forward/side scatter and cell area/height using FlowJo software. This approach was designed to exclude small particulate debris and multi-cell aggregates from the analysis.

### Donor RGC Transplantation

Donor RGC transplantations were performed at the Schepens Eye Research Institute with approval from the Institutional Animal Care and Use Committee. To support donor RGCs and establish a proneuronal microenvironment, mouse or human donor RGCs were formulated with pro-survival factors. To avoid repeated intraocular injections, which could cause additional inflammation and reduce donor RGC viability, we formulated donor RGCs with slow-release BDNF and GDNF using the Polyhedrin Delivery System (PODS Human BDNF and PODS Human GDNF (Cell Guidance Systems, PPH1-50 and PPH2-50). Donor RGCs were transplanted (10,000 – 20,000 RGCs/retina) into adult (1- to 4-month-old) RGC-deficient mouse retinas (inherited loss or neurotoxic model) with and without BDNF- and GDNF-loaded PODS under a general (ketamine/xylazine intraperitoneal injections) and local (0.5% proparacaine eye drops) anesthetic. Donor RGC suspensions (1 µL) were injected intravitally using a beveled glass microneedle with an inner diameter of 80 µm at a flow rate of 1 µL per minute. Mice were kept on a standard 12-hour light-dark cycle and were euthanized at predetermined endpoints—2 weeks for transplants from the same species (syngeneic) and 3 days for cross-species (xenotransplantation) transplants— using CO2 inhalation. Three to fourteen days after transplantation, the eyes were enucleated, fixed for 24 hours at room temperature in 4% paraformaldehyde, and then processed for retinal flat-mount preparation. Retinas were stained for host and donor cells to assess RGC survival and neurites (8 – 11 mice/group). Specifically, eyes were enucleated after three days for human donor RGC transplants to minimize immune reactions typical of xenotransplantation and after fourteen days for mouse donor RGCs to allow for neurite outgrowth.

### Electroretinogram

To measure functional recovery following transplantation, mice were injected 2 μL of 20 mM with NMDA 1 week prior to intravitreal injection of donor RGCs to cause RGC death and subsequently received either cells alone, cells+PODS, or PODS alone. Electroretinography (ERG) was performed six weeks post-transplant to assess host retinal function. Before recording electrical activity in the retina with ERG, mice were dark-adapted overnight. Mice were then anesthetized with ketamine (120 mg/kg) and xylazine (20 mg/kg), administered intraperitoneally with a 25G needle. The pupils of each mouse were dilated using 1% Tropicamide, followed by the application of Genteal to maintain corneal moisture. Mice were positioned on a temperature-controlled platform on the ERG machine (Diagnosys LLC). Reference and ground electrodes were placed under the skin on the forehead and tail, respectively. Two gold-ring electrodes were carefully positioned on the corneas without blocking the pupil. A drop of artificial tear solution was applied to ensure hydration throughout the experiment. The positive scotopic threshold response (pSTR) data were gathered using a series of flashes, averaging 40 responses for each intensity level. The pSTR was quantified by measuring the amplitude from the baseline to the peak of the positive wave. Post-ERG, the eyes were treated with antibiotic ointment and artificial tears. The mice were then placed on a warming pad with a circulation of heated water to stabilize their body temperature, allowing for a safe recovery.

### Immunohistochemistry

To visualize RGCs in culture, retinal flat-mounts, and retinal sections, samples were fixed for 30 min for cell culture and >4 hours for tissue samples with 4% (v/v) paraformaldehyde at room temperature. For retinal sections, eyes were sent to the Schepens Eye Research Institute Morphology Core for paraffin embedding (producing 6 μm thick sections). The paraffin-embedded sections were first treated with an xylene substitute, followed by ethanol washes. Antigen retrieval was then performed using a sodium citrate buffer (10 mM Sodium citrate, 0.05% Tween-20, pH 6.0) for 30 minutes.

For immunofluorescence staining, samples were incubated overnight (>12 hours) at 4°C with 1:400 goat anti-mouse secondary antibody in blocking buffer (10% goat serum, 1% bovine serum albumin, 0.1% sodium citrate, 0.25% Tween-20, 0.25% Triton-X, and 0.3M glycine in 1x phosphate-buffered saline) to reduce non-specific staining. Following blocking, the RGCs and retinal tissues were incubated with primary antibodies in staining buffer (1% bovine serum albumin, 0.25% Tween-20, and 0.25% Triton-X in 1x PBS) for 48 to 72 hours at 4°C. A complete list of primary antibodies and their concentrations is provided in Table S1. Post-primary antibody incubation, tissues were washed three times in wash buffer (0.1% Tween-20, 0.1% Triton-X in 1x PBS) to remove unbound antibodies. Secondary antibodies (goat anti-chicken, goat anti-rabbit, and goat anti-mouse) were added at a 1:500 dilution in staining buffer and incubated for 12 to 24 hours at 4°C. Following another triple wash, the samples were stained with 1 μg/mL DAPI in PBS for 20 minutes to highlight cell nuclei. To reduce tissue autofluorescence, samples were treated with 10 mM copper sulfate (pH 5.0) at room temperature for 15 minutes. After a final PBS rinse, the samples were mounted between glass cover slides and coverslips using a polyvinyl alcohol-based mounting medium containing an antifade agent (1,4-Diazabicyclo[2.2.2]octane). Imaging was performed using an Olympus IX83-FV3000 confocal microscope.

### Enzyme-linked immunosorbent assay (ELISA)

BDNF ELISA (Boster #EK0307) was performed according to the manufacturer’s protocol. PBS solution (50 µl) containing either 1e6, 5e5, 2e5, 5e4, or 1e4 PODS BDNF was added to wells of a 96-well plate and was spun down at 3000 g for 25 min. PBS was removed, and the plate was dried in a laminar flow cabinet. RPMI 1640 Complete Medium supplemented with 10% bovine calf serum was added to each well (100 µl/well). The plate was then incubated at 37 °C, 5% CO2. For the release time curve, medium from wells with 1e6 PODS was collected on days 1, 2, 3, 4, and 7 after medium addition and stored at -20 °C. For the dose-response curve, the medium was collected on day 4 after medium addition.

### Extracellular recordings

Human stem cell-derived RGC neural activity was quantified using a high-density multielectrode array (HD-MEA). HD-MEA chips (MaxOne, MaxWell Biosystems) consist of 26,400 electrodes arranged in a 120 × 220 configuration with 17.5 μm center-to-center electrode spacing to create a 3.85 × 2.10 mm^2^ sensor area.^22^ Each HD-MEA chip was cleaned using a 1%-Terg-a-zyme-solution, sterilized with 70% ethanol, and coated with 0.1 mg/mL Poly-D-Lysine (PDL) in 1x borate buffer and 0.025 mg/mL laminin in N2B27 [1x GlutaMAX, 1x antibiotic antimycotic, 1% N2 supplement, and 2% B27 supplement in 50:50 1x DMEM/F12:Neurobasal media] before use. RGCs (∼5 uL, 20,000 RGCs/uL) were seeded directly onto the sensor area of each MEA chip at a density of ∼12,500 RGCs/mm2 with or without BDNF- and GDNF-loaded PODS (150 PODS/µL) and allowed to attach in standard culture conditions for 2 hours. After attachment, 1 – 2 mL N2B27 was added to each chip (depending on the chip dimensions), and 50% of the media was replaced every other day.

After 18 – 20 days in culture, cell culture media was replaced with warmed BrainPhys^TM^ media 1 hour before extracellular recording. Spontaneous neural activity was recorded from donor RGCs using the MaxOne HD-MEA system (MaxWell Biosystems). The MaxOne HD-MEA system was placed in a cell culture incubator (37°C, >95% humidity, 5% CO2) during the recordings, and RGC activity was recorded using the MaxLab Live software (MaxWell Biosystems).

For all recordings, first, an Activity Scan Assay was performed in ‘checkered scan mode’ using a 5 – 5.5σ spike threshold to identify the active electrodes. Activity scans in ‘checkered scan mode’ provide an optimal balance between speed and resolution to identify possible neural units (RGCs with spontaneous activity). After generating an activity map, a Network Assay was performed to simultaneously record from ∼120 neural units for 5 minutes. Each neural unit consists of a 9 x 9 electrode configuration centered on active electrodes with the highest firing rate.

Spike sorting was performed on the 5-minute Network Assay recordings using the SpikeInterface framework and the SpykingCircus2 algorithm to identify individual neurons from each neural unit.^23,24^ Recordings were preprocessed using a Bandpass filter with a minimum and maximum frequency of 300 and 3000 Hz and a common mean referencing technique. Spike sorting was performed using the default parameters, except the radius was increased to 250 μm. The spike sorting output was automatically curated, and neurons were removed if they had a presence ratio less than 90% or an amplitude medium below 20 μV. RGC firing rate and spike amplitude were reported as the mean of a minimum of five MEA chips for the Control and PODS groups.

### Single-cell RNA Sequencing Data Analysis

Single-cell RNA sequencing data analysis was performed in RStudio v. 2022.07.2 (R language v. 4.2.1) using previously published datasets from the human fetal retina (FD59, FD82, FD105). Raw data was processed as previously described.^25,26^ For the stages FD82 and FD105, the analysis included both the peripheral and central retina. Seurat objects with annotated UMAP embeddings are available in the source data file.

To show the changes of gene expression for different timepoints, the AverageExpression function in Seurat was applied to the whole datasets. Trends for GDNF and BDNF average expression at different time points plots were visualized using ggplot2 (v. 3.3.6) and ggalt (v. 0.4.0) packages. The analysis also included a step with correlation between NTRK2, GDNF receptor, and GFRA1, GDNF receptor and apoptosis pathway genes expression. To obtain the gene list for apoptosis, we downloaded the annotation from the Molecular Signatures Database (http://www.gsea22msigdb.org/gsea/msigdb/human/geneset/HALLMARK_APOPTOSIS.html).

To check the apoptosis pathway expression, we used the escape package 1 (v. 1.6.0) (12). We merged subsetted RGCs populations in FD59, FD82, and FD125 Seurat objects and performed the enrichment on the RNA count data using the enrichIt function with the ssGSEA method (13). To examine the correlation between NTRK2/GFRA1 and the apoptosis pathway expression, we generated a plot using FeatureScatter Seurat function for the GFRA1 positive and NTRK2 positive cells. We added regression model to the analysis using geom_smooth function with ‘lm’ method. The software and packages used for the analysis can be found in Table S2. The code used for this study to reproduce the bioinformatic analysis is available on GitHub at https://github.com/mcrewcow/BaranovLab/blob/main/survival_RGC_paper.R.

### Statistical Analysis

Statistical significance was calculated using GraphPad PRISM 10 using an unpaired t-test or a Tukey one-way ANOVA. Error bars represent the mean ± SD of measurements (*P < 0.05, **P < 0.01, ***P < 0.001, and ****P < 0.0001). Each dot within a bar plot represents an individual retina/mouse or well/MEA.

## Results

### BDNF and GDNF Treatment Improves RGC Structural and Function Maturation *In Vitro*

Growth factors have been widely described for supporting neuron survival, differentiation, and neurite outgrowth.^27,28^ To evaluate the impact of BDNF and GDNF on retinal ganglion cell (RGC) differentiation and function, we utilized mouse and human stem cell-derived retinal organoid models. Mouse RGCs were differentiated from a Thy1.2-GFP iPSC reporter line, as previously described, allowing visualization and quantification of RGC yield via GFP expression (**Figure 1A-B**). After three weeks of differentiation, the cultures were supplemented with a slow-release formation of BDNF and GDNF (BDNF and GDNF-loaded polyhedrin-based particles (PODS, Polyhedrin Delivery System). BDNF and GDNF-loaded PODS show approximately 50 pg/mL sustained growth factor release in culture over a week (**Figure S1**). Flow cytometry analysis of dissociated organoids at day 27 revealed a significant increase (>50%) in Thy1.2-positive cell populations in cultures treated with BDNF/GDNF-PODS compared to untreated controls **(Figure 1B-C; Figure S2**).

**Figure 1.**
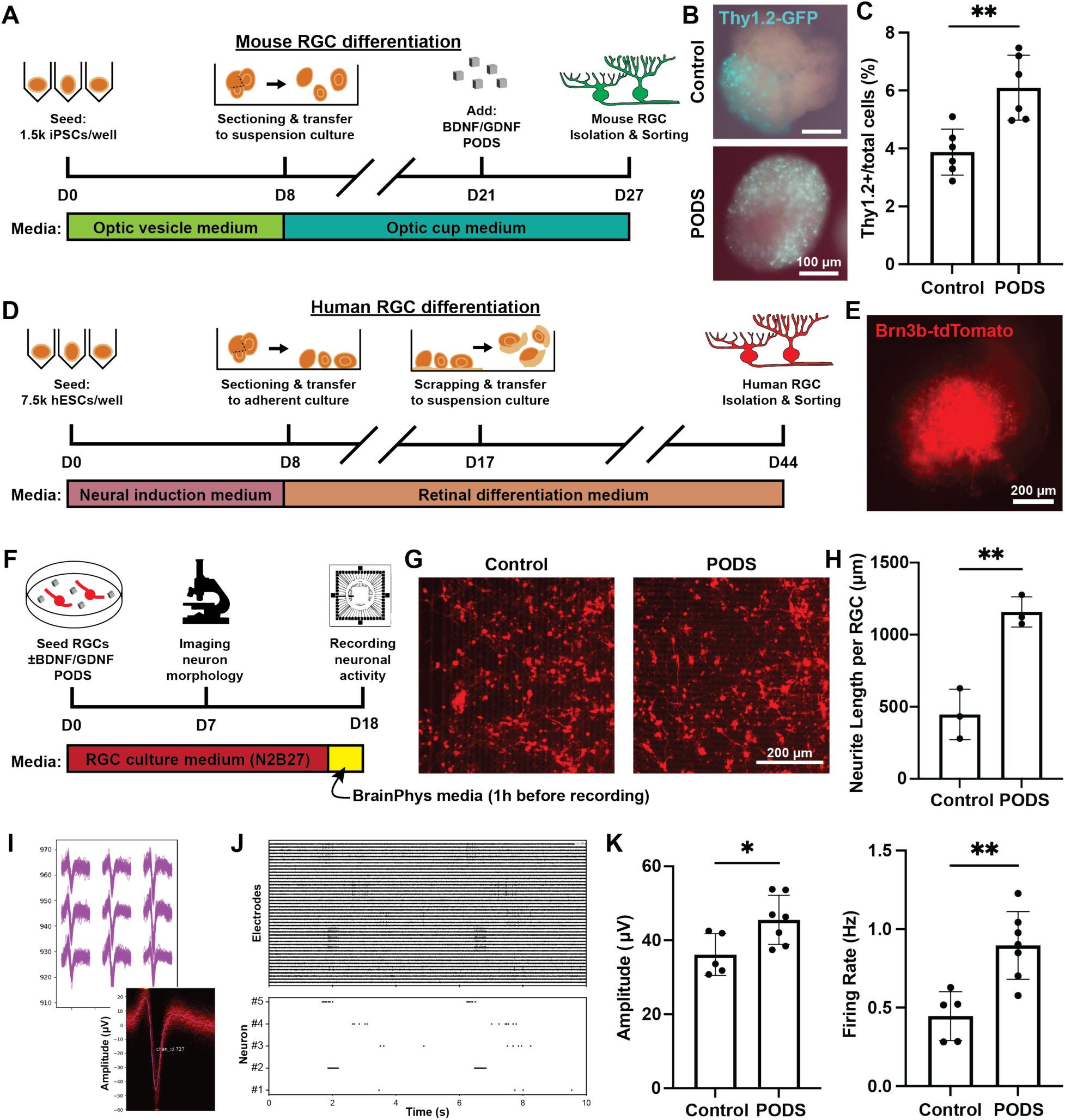
Human stem cell-derived RGC culture in response to BDNF/GDNF-PODS co-treatment. **(A)** Schematic overview summarizing the differentiation of mouse Thy1.2-GFP iPSCs to retinal organoids. Organoids were sectioned on day 8 to improve nutrient transport and limit cell death. Retinal organoids were differentiated over 3 weeks before being supplemented with BDNF/GDNF-PODS for 1 week. **(B)** Representative fluorescent image of late-stage (D27) mouse retinal organoids with and without PODS co-treatment during the differentiation. **(C)** Quantification of Thy1.2+ yield by flow cytometry in response to PODS co-treatment. **(D)** Schematic overview summarizing the differentiation of human BRN3B-tdTomato ESCs to retinal organoids. Organoids were sectioned on day 8 to improve nutrient transport and limit cell death, and they were transitioned between 2D and 3D cultures to better recapitulate human development. **(E)** Representative fluorescent image of human retinal organoid after 44 days of differentiation shows robust tdTomato expression. **(F)** Schematic overview summarizing the timeline to evaluate the effect of BDNF/GDNF-PODS on human RGC neurite outgrowth and neural activity. **(G)** Representative fluorescent images of td-Tomato positive RGCs cultured on an MEA with and without BDNF/GDNF-PODS. **(H)** Quantification of neurite outgrowth after 1 week in culture. **(I)** Representative spontaneous RGC action potentials recorded from multiple electrodes corresponding to a single neuron using a high-density multielectrode array. **(J)** Representative traces from preprocessing MEA recordings of spontaneous RGC action potentials from individual electrodes after 18 days in culture and accompanying raster plot showing five neurons identified after spike sorting the raw traces. **(K)** Quantification of median amplitude and mean spontaneous firing rate measured from all the individual neurons cultured in an MEA with and without BDNF/GDNF-PODS.

In parallel, human stem cell-derived RGCs were generated using the 3D retinal organoid model to confirm that BDNF/GDNF-PODS would improve RGC neurite outgrowth. Human RGCs were differentiated using a Brn3b-tdTomato human stem cell reporter line to visualize RGCs with tdTomato (**Figure 1D-E**), as previously described. To characterize the effect of BDNF and GDNF on these stem cell-derived RGCs, we evaluated neurite outgrowth in vitro 1 week after adding BDNF/GDNF-PODS (**Figure 1F**). Cultures supplemented with BDNF/GDNF-PODS showed significantly enhanced neurite outgrowth, with a mean neurite length of 1053 ± 105.3 µm, compared to 517 ± 105.3 µm in untreated controls (p<0.01; **Figure 1G-H**).

Lastly, to evaluate the functional impact of BDNF/GDNF-PODS, human RGCs were cultured on high-density multielectrode arrays (MEAs) to record spontaneous action potentials, an analog for functional maturation.^29^ Human stem cell-derived RGCs were seeded on an MEA with and without PODS and cultured for three weeks. Due to the high-density nature of the MEA, individual RGC actional potentials were recorded from multiple electrodes simultaneously (**Figure 1I**). To resolve individual neurons, we performed spike sorting before analyzing the data (**Figure 1J**). We detected no spontaneous action potentials within the first week of culture; however, by 18 – 20 days, human donor RGC exhibited a robust firing response. Those RGCs cultured with PODS significantly increased their median spike amplitude and mean firing rate compared to controls. PODS co-treatment resulted in a 45.5 µV median spike amplitude and 0.90 Hz mean firing rate compared to 36.1 µV and 0.44 Hz in the control (**Figure 3D-E**).

### BDNF and GDNF are Key Neurotrophic Factors in Retinal Development and RGC Survival

To further explore the role of neurotrophic factors in retinal ganglion cell (RGC) survival and retinal development, we performed an in silico analysis of single-cell transcriptomic data from developing (FD59, FD82, and FD125) and adult human retinas.^30,31^ Our analysis revealed that both BDNF and GDNF were highly expressed during early developmental stages, corresponding precisely with critical periods of RGC differentiation and maturation (**Figure 2A**). Notably, the expression of these factors decreased significantly as the retina matured, suggesting that reintroducing BDNF and GDNF into the retinal microenvironment may restore conditions favorable for RGC regeneration and repopulation.

**Figure 2.**
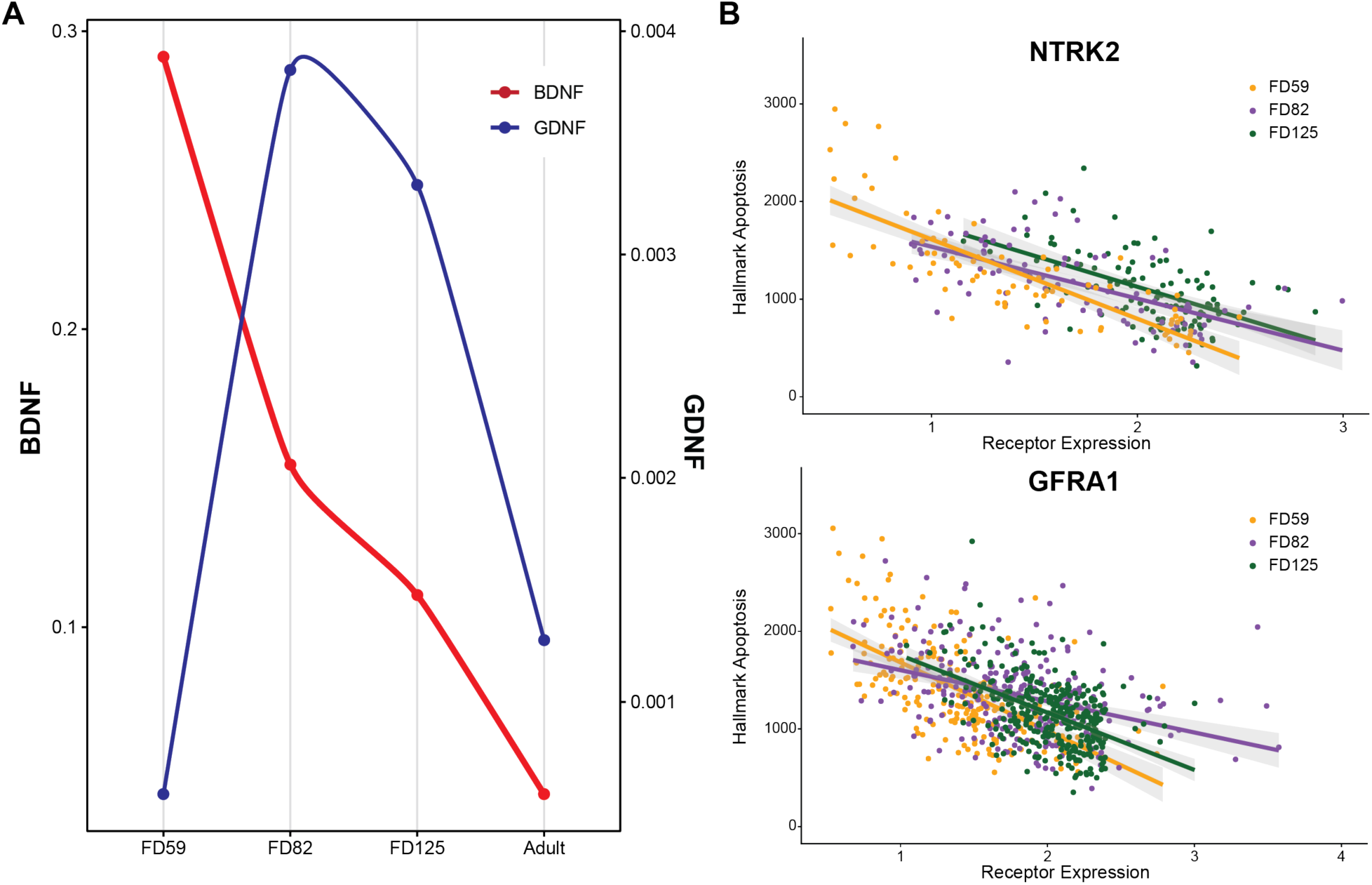
Characterization of neurotropic factors in the retinal microenvironment during development. **(A)** Expression levels of BDNF and GDNF in the retina during different stages of development. **(B)** Correlation between BDNF and GDNF receptor expression and apoptosis pathways enrichment score.

Furthermore, we investigated the correlation between BDNF and GDNF receptor expression and genes associated with apoptosis pathways. Our analysis demonstrated a robust negative correlation (**Figure 2B-C**), indicating that higher expression levels of these neurotrophic factors and their receptors were linked to decreased activation of apoptosis-related genes. This inverse relationship underscores the protective role of BDNF and GDNF in RGC survival and highlights their therapeutic potential for supporting donor RGC survival in the hostile glaucomatous retinal microenvironment.

### Modifying the Retinal Microenvironment with Neurotrophic Factors Improves Donor and Host Neuron Survival

We assessed whether supplementing the retinal microenvironment with BDNF and GDNF could enhance the survival of transplanted donor RGCs. To maintain sustained availability and mitigate rapid clearance, we continued to utilize a slow-release delivery system (BDNF/GDNF-PODS). Using mouse models of RGC loss (NMDA-induced toxicity and an inherited loss model) to mimic glaucoma (**Figure S3**), we transplanted stem cell-derived mouse RGCs in the presence or absence of BDNF/GDNF-PODS. PODS co-treatment significantly improved transplantation outcomes, increasing the transplantation success rate, defined as a transplant with over 0.5% survival of donor RGCs,^5^ from 37% to 73%. Additionally, PODS enhanced donor RGC coverage across the retina by 2.7-fold (**Figure 3A-B**). We also observed extensive neurite outgrowth toward and into the optic nerve head (**Figure 3C-D**). Although neurite complexity was not quantitatively assessed in vivo, in vitro findings strongly suggest a direct role for BDNF/GDNF-PODS in enhancing neurite extension (**Figure 1H**). This finding is critical as it demonstrates that the transplanted cells have the potential to connect with their postsynaptic targets in the brain, a necessary step for functional recovery after injury.

**Figure 3.**
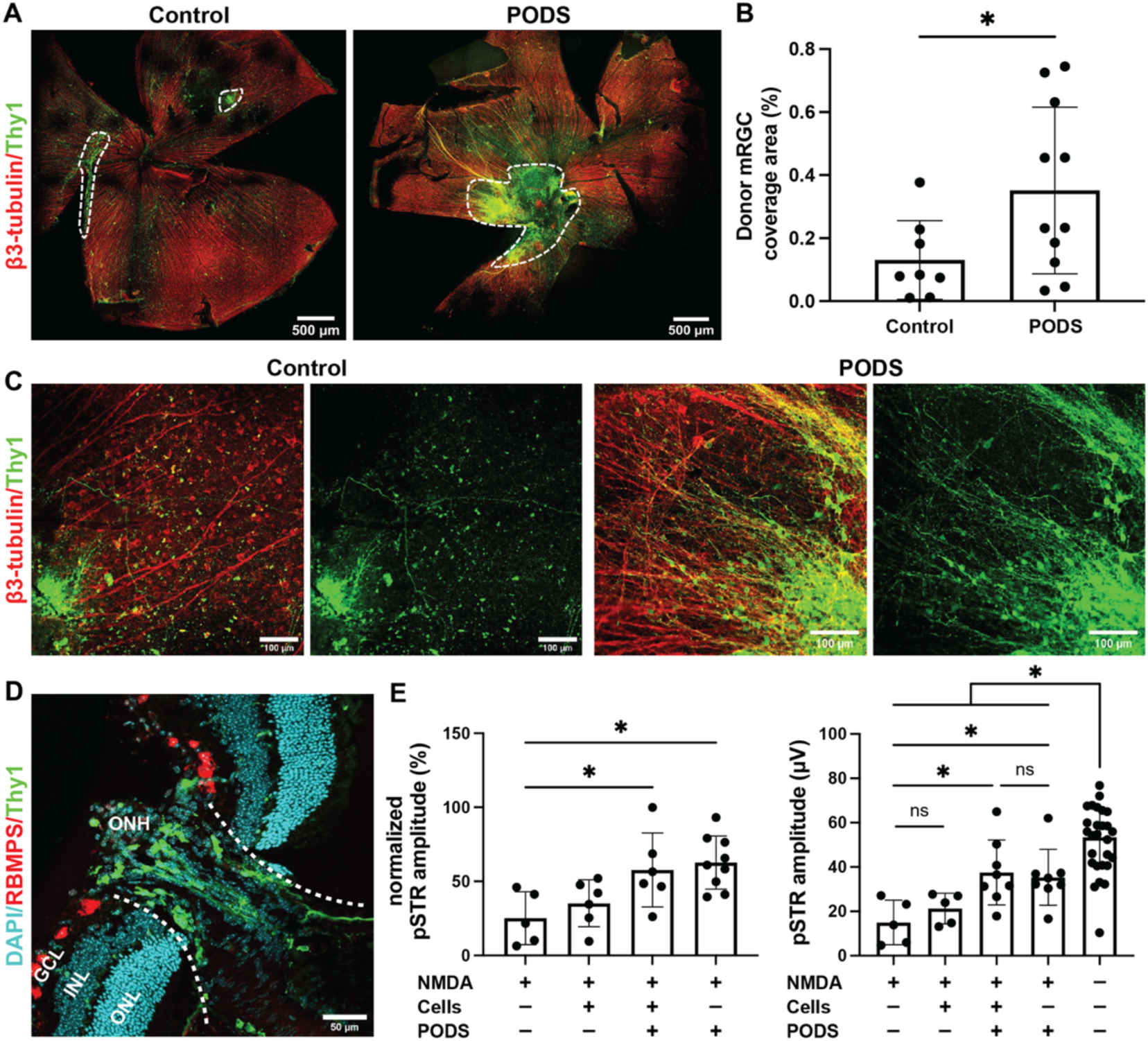
Mouse stem cell-derived RGC syngeneic transplantation in response to BDNF/GDNF co-treatment. **(A)** Representative retinal whole mount showing mouse donor RGC outgrowth (outlined) and **(B)** quantification of coverage area 2 weeks post-transplantation. **(C)** High-magnification images showing mouse donor RGC neurite extensive outgrowth with BDNF/GDNF-PODS co-treatment. **(D)** Immunofluorescent image of a retinal section showing donor RGC entry into the optic nerve head (ONH) and down the optic nerve (outlines). GCL, ganglion cell layer; INL, inner nuclear layer; ONL, outer nuclear layer. **(E)** Quantification of positive scotopic threshold response (pSTR) measurements after transplantation and BDNF/GDNF-PODS co-treatment shows an improvement in host RGC function in an NMDA-induced toxicity model as measured by electroretinogram (ERG).

To evaluate retinal function after transplantation, we measure the positive scotopic threshold response using electroretinography (ERG) to evaluate RGCs.^32,33^ While our mouse stem cell-derived RGCs were shown to survive and integrate within the host retina (**Figure 3A-D**), our ERG results indicate that Thy1-GFP+ donor RGCs alone don’t rescue retinal function at six weeks post-transplantation in the NMDA model of toxic optic neuropathy (**Figure 3E**). However, treatment with PODS alone or with cells+PODS led to the partial preservation of RGC function as measured by pSTR response at six weeks (32uV in the PODS group vs. 17uV in the control group, **Figure 3E**). This indicates a neuroprotective effect of BDNF/GDNF-PODS on host RGCs.

In addition to mouse stem cell-derived RGC transplantation, to improve the clinical relevance of this work, we transplanted human stem cell-derived RGCs into mouse retinas. To mitigate immune responses associated with xenotransplantation, retinas were analyzed 3 days post-transplantation. Although the PODS co-treatment did not affect the success rate of human donor RGC transplantation quantified by the number of animals that had donor RGCs with neurites in 3 days, the co-delivery of slow-release formulation led to a 15-fold increase in the total number of human donor RGCs (**Figure 4A-C**). Furthermore, despite only 3 days of growth post-transplantation, 50% of PODS-treated retinas displayed donor RGC neurites extending toward the optic nerve head, with a significantly greater average length (1008.44 ± 264.12 µm) compared to only 10% in untreated controls (**Figure 4D-E**). These results demonstrate that sustained-release BDNF/GDNF formulations improve the survival and integration of transplanted human RGCs, highlighting their potential for supporting functional recovery in retinal degenerative diseases.

**Figure 4.**
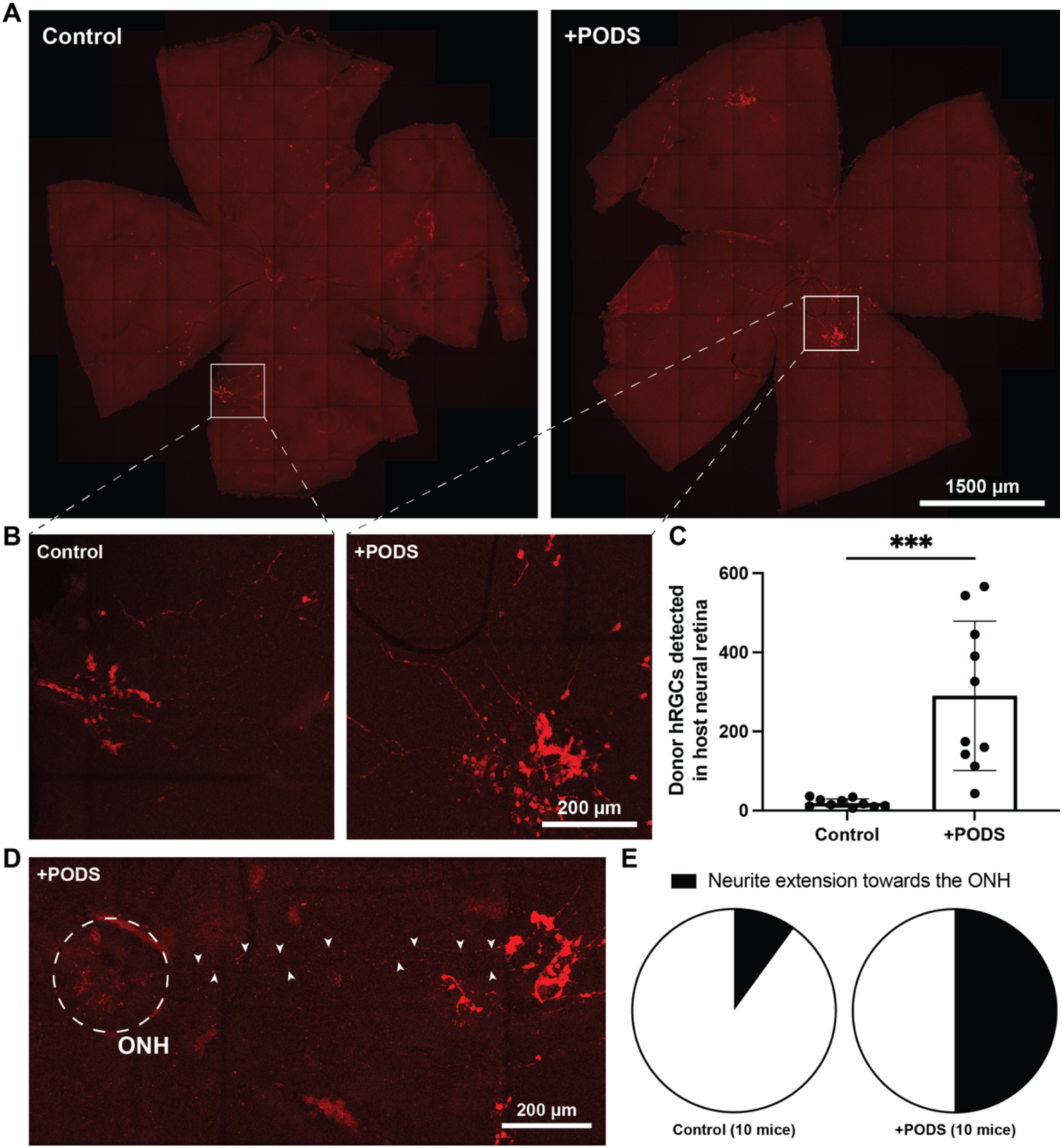
Human stem cell-derived RGC xenotransplantation in mice supplemented with sustained-release neurotrophic factors. **(A)** Representative retinal flat mounts and **(B)** high-magnification images showing human donor RGC outgrowth (red). **(C)** Quantification of the number of donor RGCs detected in the host retina for RGCs formulated with and without PODS. **(D)** Representative image of a retinal flat mount that highlights donor RGC neurite extension toward and into the optic nerve head (ONH). White arrows point to neurites extending the ONH. **(E)** Quantification of the number of transplanted retinas showing neurite extension toward the ONH with and without PODS co-treatment 3 days post-transplantation.

## Discussion

Blindness and vision impairments resulting from the loss or degeneration of RGCs are irreversible. While the underlying pathology of optic neuropathies is diverse (e.g., elevation of intraocular pressure, acute trauma), all phenotypically resemble neurodegenerative disease.^34^ Patients initially experience decreased visual fields, dyschromatopsia, and altered pupillary responses, gradually progressing toward complete vision loss.^35^ Optical coherence tomography and other live imaging approaches have shown slow RGC death in glaucoma,^36^ underscoring the importance of timely intervention. Applying pro-survival factors early in disease progression may preserve remaining function and partially reverse vision loss.^37–39^

In this study, we demonstrated that sustained delivery of BDNF and GDNF significantly enhances RGC survival, differentiation, and integration following transplantation. Both BDNF and GDNF are well-established as critical factors in retinal development and neuronal survival. Consistent with our findings, Fligor et al. also reported increased RGC numbers following BDNF treatment, as visualized by a BRN3B reporter in human retinal organoids.^40^

Prior studies support the beneficial effects of BDNF and GDNF on RGC survival and regeneration both in vitro and in vivo. In vitro studies have shown that BDNF robustly increases neurite length and branching in cultured rat RGCs.^41^ Additionally, Meyer-Franke et al. demonstrated that BDNF significantly improves the survival of rat RGCs in culture, emphasizing its critical role in neuronal maintenance.^42^ Despite these promising results, translating neurotrophic factors into clinical applications has faced challenges, including rapid clearance and limited bioavailability within the ocular environment.^43,44^ For instance, intravitreal injections of BDNF combined with a free radical scavenger delayed RGC apoptosis and preserved visual function in rodent glaucoma models but required frequent injections.^45^ To address this limitation, slow-release formulation can be designed, or gene therapies can be used to maintain sustained availability. Notably, prolonged BDNF delivery through gene therapy facilitated axon regeneration and partial vision restoration following an optic nerve injury in adult rats.^46^

Similarly, GDNF-loaded microspheres have shown neuroprotective properties in preclinical models of glaucoma and ischemic retinal injury. Intravitreal administration of GDNF significantly reduced RGC loss and preserved visual function in rodent glaucoma models.^47^ Furthermore, recombinant adeno-associated virus-mediated delivery of GDNF also enhanced RGC survival and reduced optic nerve degeneration in experimental optic neuropathies.^48^

Our transcriptomic analyses further support the therapeutic value of BDNF and GDNF, revealing significantly reduced expression in the adult retina compared to developmental stages. This observation underscores the importance of maintaining adequate levels of BDNF and GDNF in the diseased retinal environment, where natural expression may decline significantly following RGC loss, further exacerbating the degenerative process. Supplementation of BDNF and GDNF thus provides dual benefits, directly supporting transplanted RGCs and offering neuroprotection to the host retinal tissue. Further exploration of developmental retinal transcriptomics may uncover additional factors beneficial for transplantation and neuroprotection, complementing traditional discovery approaches such as phenotypic screenings and loss-of-function animal models.^49–51^

Although neurotrophic factors can support host neuron survival, once RGCs are lost, these factors alone cannot restore them. Stem cell technologies offer new avenues for cell replacement therapies. Clinical trials and research studies have shown the potential of retinal pigment epithelium and photoreceptor replacement to halt visual decline in age-related macular degeneration and cone/rod dystrophies.^52,53^ Similarly, glaucoma, a progressive neurodegenerative disease, could benefit from cell replacement to prevent or recover vision loss ^6^. However, replacing RGCs is particularly challenging because donor neurons need to both structurally and functionally integrate within the host retina and extend axons toward the optic nerve and brain.

RGC replacement by transplantation of primary or stem cell-derived RGCs is a potential approach to enable vision recovery within advanced-stage optic neuropathies.^6^ Our previous work showed that over half of healthy and even RGC-depleted recipient retinas retained donor RGCs post-transplant.^5^ However, the number of surviving donor RGCs and retinal area covered by the transplant remained low. To overcome this, we co-delivered BDNF and GDNF using the slow-release PODS system, creating a more favorable retinal microenvironment for regeneration. Our current study demonstrates that this strategy not only increases donor RGC survival but also promotes neurite growth toward the optic nerve head (**Figures 3 and 5**). This approach likely supports transplanted cells during their crucial initial integration period, reducing apoptosis and enhancing integration.

Electroretinography assessments further demonstrated broader neuroprotective benefits, showing partial preservation of host retinal function through sustained neurotrophic factor delivery. Thus, our approach not only enhances donor cell survival but also provides neuroprotection to host retinal neurons, ultimately reducing overall neurodegeneration.

## Conclusions

Modulating the host retinal microenvironment with sustained-release growth factors represents an innovative approach to improving transplantation outcomes. Furthermore, enhancing donor RGC morphology and axon outgrowth opens new possibilities in regenerative medicine for treating glaucoma and other optic neuropathies. Using neuroprotective factors in combination with other approaches to modify the retinal microenvironment and stem cell-derived RGC delivery may offer a viable solution to the challenge of RGC replacement. Depending on the model or state of the disease, the retinal microenvironment may also need to be altered to support donor RGC migration, growth, maturation, axon extension, and increased metabolic demands.^6,19^ Integrating neuroprotective strategies with cell replacement therapies is a promising direction for research and clinical applications. Lastly, sustained-release growth factor formulations, such as PODS, represent a promising neuroprotective approach to limit RGC death before loss. Altogether, these findings provide promising insights into potential strategies for restoring vision lost due to RGC damage or loss in optic neuropathies.

## Description of the Supporting Information Material

Figure S1 to S3

Tables S1 to S2

## Conflict of Interest Disclosure

The Schepens Eye Research Institute and Cell Guidance Systems have a patent incorporating the polyhedrin-based slow-release growth factor system described in this report with inventors J.O., C.P., and P.B.

## Author Contributions

Conceptualization, JRS, JO, PB; Methodology, JRS, EK, JO, PB; Software: JRS, EK; Validation, JRS, EK, PB; Formal analysis, JRS, EK, PB; Investigation, JRS, EK, JO, MP, JM, CP, PB; Resources, MP, CP, PB ; Data Curation, JRS, EK; Writing - Original Draft, JRS; Writing - Review & Editing, JRS, EK, JO, MP, JM, CP, PB; Visualization, JRS, EK; Supervision, PB; Project administration, PB; Funding acquisition, CP, JRS, PB

## Acknowledgment

National Institutes of Health/National Eye Institute grant K99EY036122 (JRS); National Institutes of Health/National Eye Institute grant F32EY033211 (JRS); National Institutes of Health/National Eye Institute award L70EY034355 (JRS); National Institutes of Health/National Eye Institute grant U24EY029893 (PB); National Institutes of Health/National Eye Institute grant P30EY003790 (SERI Morphology Core); Bright Focus Foundation grant G2020231 (PB); Gilbert Family Foundation Vision Research Initiative grant (PB); Department of Defense VRP FTTSA (PB).

## Abbreviations

RGC: retinal ganglion cell
PODS: Polyhedrin Delivery System
BDNF: brain-derived neurotrophic factor
GDNF: glial-derived neurotrophic factor
HD-MEA: high-density multielectrode array
pSTR: positive scotopic threshold response
ONH: optic nerve head
ERG: Electroretinography
ssGSEA: single-sample gene set enrichment analysis.

## Supporting Information

**Figure S1.**
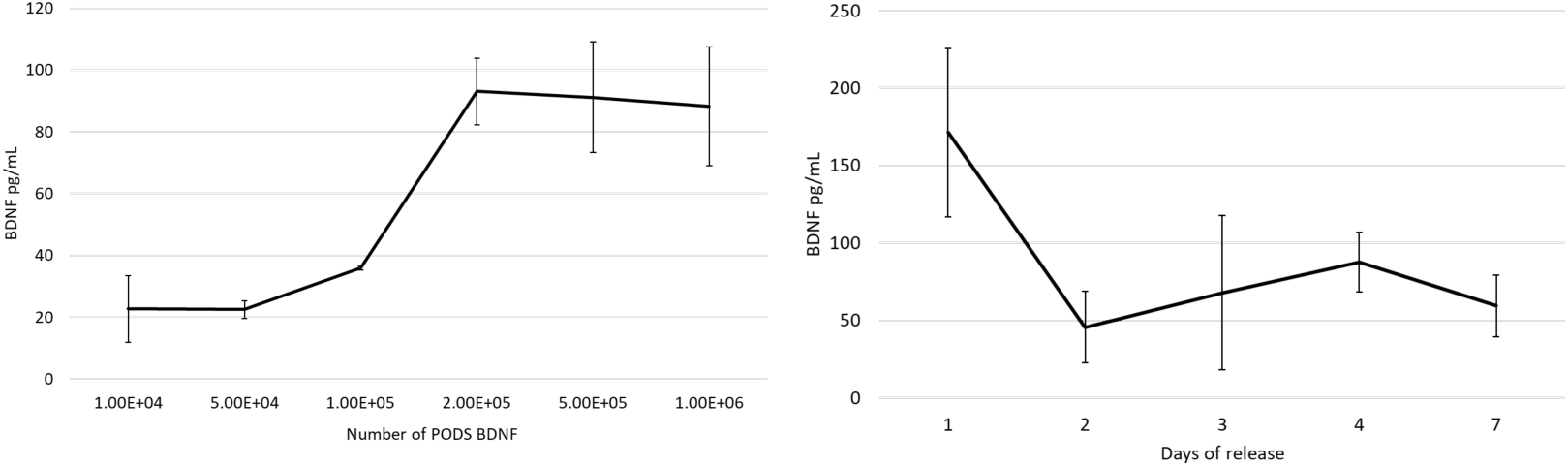
Release Kinetics of BDNF from PODS crystals. The availability of BDNF in the cell culture medium, released from PODS Human BDNF, is both time- and dose-dependent.

**Figure S2.**
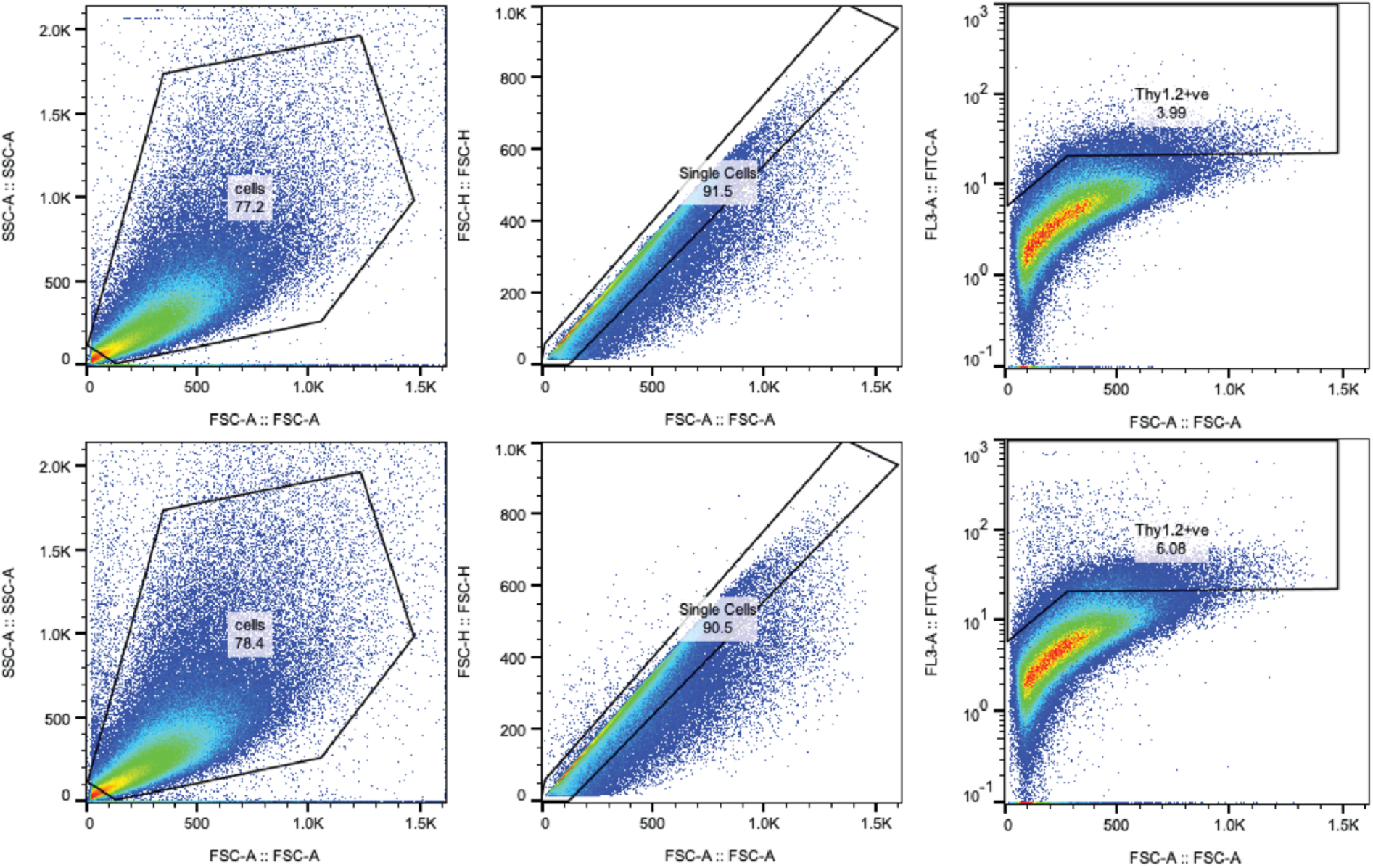
Slow-release BNDF and GDNF improve RGC differentiation yield. Representative FACS plots with gating for dissociated mouse retinal organoids in the control group (top) vs. the PODS group (bottom).

**Figure S3.**
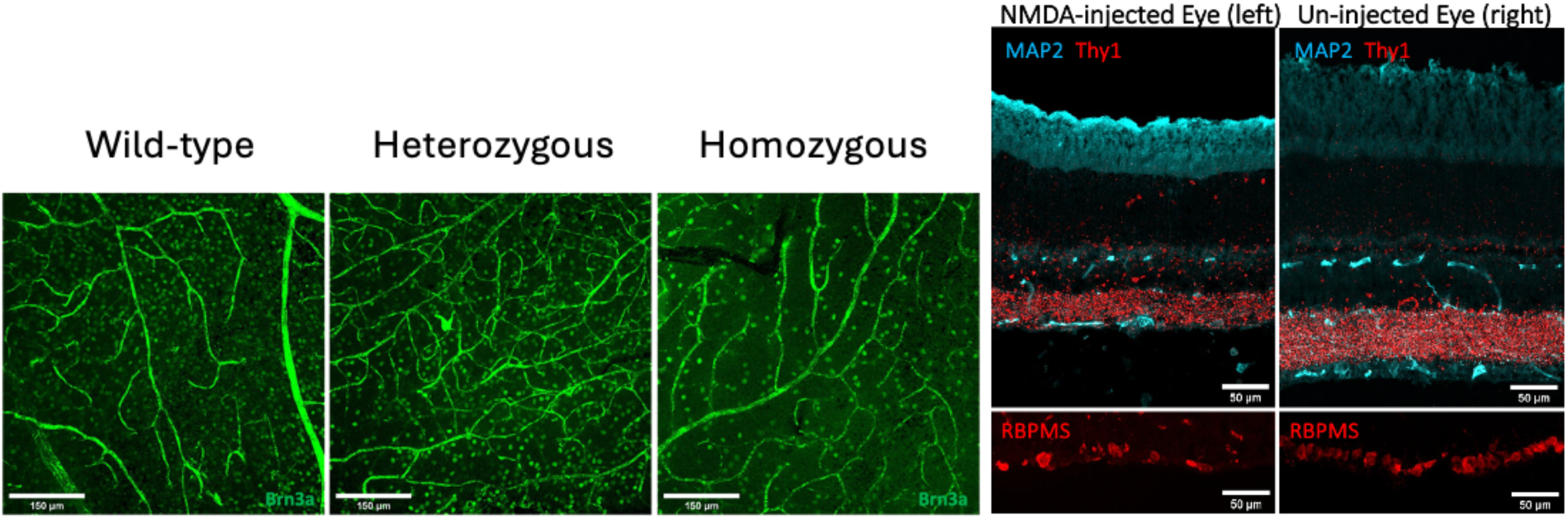
Inherited loss or neurotoxic mouse models. Representative fluorescent images of retinal flat mounts and sections showing RGC loss in an inherited loss model (homozygous) and neurotoxic models (NMDA-injected).

**Table S1.** Primary antibodies for immunohistochemistry.

| **Antibody** | **Concentration** | **Vendor** | **Catalog #** |
| --- | --- | --- | --- |
| Chicken Anti-mCherry | 1:500 | Abcam | ab205402 |
| Chicken Anti-GFP | 1:50 | enQuire<br>BioReagents | NC1666473 |
| Rabbit Anti-RBPMS | 1:300 | GeneTex | GTX118619 |
| Mouse Anti-Thy1 | 1:400 | Serotec | MCA02R |
| Rabbit Anti-Map2 | 1:250 | Biolegend | 840601 |
| Mouse B3-tubulin | 1:500 | Sigma Aldrich | T8660 |
| Mouse Brn3a | 1:500 | Millipore | MAB1585 |

**Table S2.** Software packages for single-cell RNA sequencing analysis.

| Software<br>/package | Version | Link |
| --- | --- | --- |
| R | 4.2.1-4.3.1 | https://cran.r-project.org/bin/ |
| RStudio | 2022.07.<br>2-2023.06.<br>1 | https://www.rstudio.com/products/rstudio/download/ |
| Azimuth | 0.4.6 | https://github.com/satijalab/azimuth |
| biomaRt | 2.52.0-2.56.1 | https://bioconductor.org/packages/release/bioc/html/biomaRt.ht<br>ml |
| CellChat | 1.5.0-2.1.0 | https://github.com/sqjin/CellChat |
| ComplexHeat<br>map | 2.16.0 | https://bioconductor.org/packages/release/bioc/html/ComplexHe<br>atmap.html |
| cowplot | 1.1.1 | https://cran.r-project.org/web/packages/cowplot/ |
| dittoSeq | 1.8.1-1.12.2 | https://bioconductor.org/packages/release/bioc/html/dittoSeq.htm<br>l |
| dplyr | 1.0.10-1.1.3 | https://dplyr.tidyverse.org/ |
| escape | 1.5.1-1.6.0 | https://github.com/ncborcherding/escape |
| future | 1.33.0 | https://cran.r-project.org/web/packages/future/index.html |
| ggalt | 0.4.0 | https://yonicd.github.io/ggalt/index.html |
| ggplot2 | 3.3.6-3.4.3 | https://ggplot2.tidyverse.org/ |
| GSEABase | 1.58.0-1.62.0 | https://bioconductor.org/packages/release/bioc/html/GSEABase.html |
| htmlwidgets | 1.5.4-1.6.2 | https://www.htmlwidgets.org/ |
| LRLoop | 0.1.0 | https://github.com/Pinlyu3/LRLoop |
| Matrix | 1.5-1-1.6-0 | https://cran.r-project.org/web/packages/Matrix/index.html |
| patchwork | 1.1.2-1.1.3 | https://patchwork.data-imaginist.com/ |
| plotly | 4.10.0 | https://plotly.com/r/ |
| reticulate | 1.26-1.32.0 | https://rstudio.github.io/reticulate/ |
| SCopeLoomR | 0.13.0 | https://github.com/aertslab/SCopeLoomR |
| sctransform | 0.4.0 | https://github.com/satijalab/sctransform |
| Seurat | 4.2.0-4.3.1 | https://satijalab.org/seurat/ |
| SeuratDisk | 0.0.0.90<br>20 | https://github.com/mojaveazure/seurat-disk |
| Python | 3.7.4 | https://www.python.org/ |
| anndata | 0.8.0 | https://anndata.readthedocs.io/ |
| loompy | 3.0.7 | https://linnarssonlab.org/loompy/ |
| matplotlib | 3.5.3 | https://matplotlib.org/ |
| numpy | 1.21.6 | https://numpy.org/ |
| palantir | 1.2 | https://pypi.org/project/palantir/ |
| pandas | 1.3.5 | https://pandas.pydata.org/ |
| pySCENIC | 0.12.1 | https://pyscenic.readthedocs.io/ |
| scanpy | 1.9.3 | https://scanpy.readthedocs.io/ |
| scFates | 1.0.6 | https://scfates.readthedocs.io/en/latest/ |
| seaborn | 0.12.2 | https://seaborn.pydata.org/ |

## Notes

### Summary of Updates

Updated the figures and text throughout to tell a clearer story.

